# NeuronMotif: Deciphering transcriptional cis-regulatory codes from deep neural networks

**DOI:** 10.1101/2021.02.10.430606

**Authors:** Zheng Wei, Kui Hua, Lei Wei, Shining Ma, Rui Jiang, Yanda Li, Wing Hung Wong, Xiaowo Wang

## Abstract

Discovering DNA regulatory sequence motifs and their relative positions are vital to understand the mechanisms of gene expression regulation. Such complicated motif grammars are difficult to be summarized from shallow models. Although Deep Convolutional Neural Network (DCNN) achieved great success in annotating cis-regulatory elements, few combinatorial motif grammars have been accurately interpreted due to the mixed signal in DCNN. To address this problem, we proposed NeuronMotif, a general backward decoupling algorithm, to reveal the homo-/hetero-typic motif combinations and arrangements embedded in convolutional neurons. We applied NeuronMotif on several widely-used DCNN models. Many uncovered motif grammars of deep convolutional neurons are supported by literature or ATAC-seq footprinting. We further diagnosed the sick neurons that are sensitive to adversarial noises, which can guide DCNN architecture optimization for better prediction performance and motif feature extraction. Overall, NeuronMotif enables decoding cis-regulatory codes from deep convolutional neurons and understanding DCNN from a novel perspective.

## Introduction

The DNA sequence is a language of life^1^. To understand life processes, it is essential to decode the grammar of DNA. One of the most important problems is deciphering transcriptional cis-regulatory code from functional DNA sequence. Deep sequencing techniques such as ChIP-seq, ATAC-seq^2^, etc. have been developed to discover the sequences with specific function or characteristic like Transcription Factor Binding Sites (TFBSs), Histone-marks (HMs) and chromatin openness. But the logic of the sequence is difficult to summarize directly. With the development of deep learning techniques, a growing number of researchers resort to the deep convolutional neural network (DCNN) for its significant advantage including automatic extraction of sequence motif (Fig. 1d) and higher prediction accuracy^3^. For example, DeepSEA^4^ and Basset^5^ model successfully use DNA sequence to predict chromatin-profiling data including TFBSs, HMs profiles and DNase I sensitivity. Among these common functions, in cis-regulatory modules, Transcription Factors (TF) regulate gene expression through binding or cobinding to specific preferred DNA sequences that occur at particular genome positions^6^. Accurately characterizing TF binding specificities and interpreting the relative positions of TFs from DCNN are vital to understand the logic of gene regulation (Fig. 1a). Unfortunately, DCNN is a black-box that is difficult to be interpreted what motif glossary or even motif grammar it exactly learns.

**Fig. 1.**
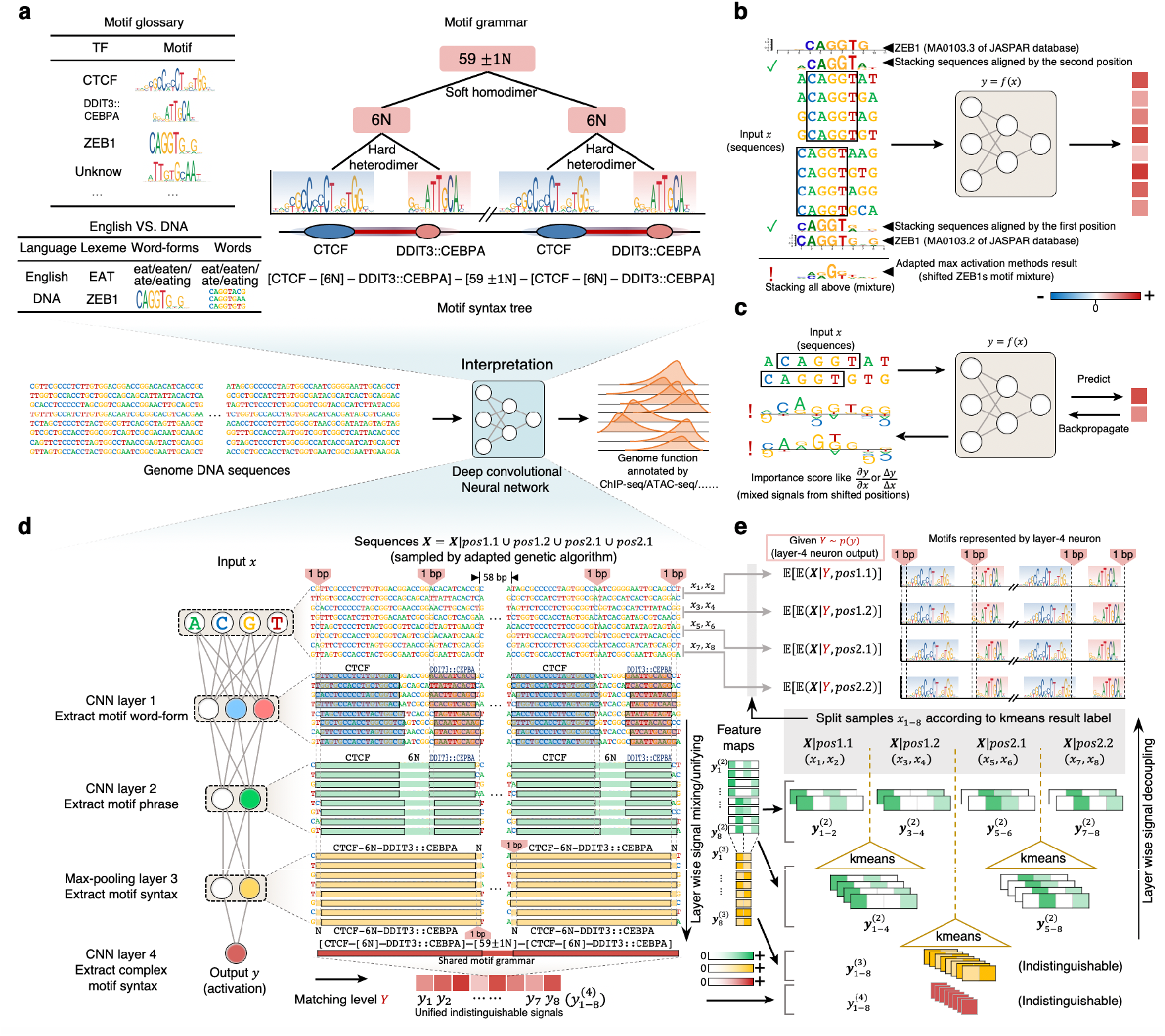
The overview of NeuronMotif and existing methods. **a**, A trained DCNN model can annotate genome function with corresponding genome sequence as input. Interpreting regulatory grammar from DCNN includes discovering the motif glossary and syntax. The motif is similar to the different word-forms for a lexeme, the smallest isolatable meaningful unit. Soft/hard hetero/homo-multimer motif are organized by motif syntax tree. **b,c**, Max activation and saliency map methods adapted from CV. **d,e** How NeuronMotif decouple a layer-4 neuron based on the mechanism of DCNN. **d**, Eight sequences ***x***_1-8_ matched by two CTCF-6N-DDIT3::CEBPA motifs with four different relative positions are sampled by adapted genetic algorithm. In each layer, the masked subsequences are detected by the neurons of the corresponding colors. Convolutional neuron combines the motif sequence recognized by previous layers (rectangles with black border) and fills the gap between them. Max-pooling operation aligns the recognized regions by extending their length. The chaotic signal of nucleotide bases in ***x***_1-8_ with similar function are layer-wisely unified into the similar signal 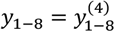 (feature map ***y***^(*l*)^, only the key components of feature map in layers *l* = 2,3,4 are shown in figure). 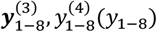 are independent of different motif sequences and shift diversity. **e**, From layer 4 to 1, feature maps of the sequences can be firstly distinguished at layer 2. To reverse the max-pooling operation of size 2, twice kmeans (k=2) are applied on feature maps 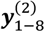 reclusively. ***x***_1-8_ are divided into 4 groups for calculating PPM respectively. *A* is the max activation in each group.

Interpretation of DCNN black-box is not as smooth as function annotations. Most existing methods^4,5,7,8^ seek to interpret DCNN by detecting the correlation between the predicted genome function as the model output and DNA sequence at the resolution of a single nucleotide base as inputs via different approaches adapted from Computer Vision (CV) (Fig. 1b, Fig. 1c and Fig S1, see supplementary information for details). However, from the viewport of linguistics, letters of nucleotide bases do not have actual meanings unless they are combined into various words of motif sequences^9^. Thus, interpreting the meaning of a single nucleotide base while ignoring its the context-dependence is polysemous or even meaningless. The average interpretation of polysemous results is a confusing mixture.

Due to the lack of interpretation methods, the design of deeper DCNN structure with better prediction performance is limited. Different from the deepening DCNNs applied in the CV like 16-convolutional-layer VGG-19 and 128-convolutional-layer ResNet^10^, most DCNN models for studying genome functions contain up to 3 convolutional layers to guarantee clear interpretations ^11,12^. The interpretation of the first layer in shallow 3-convolutional-layer DCNN is more reliable with existing interpretation methods. These shallow model avoids serious motif mixing problems in deeper layers^13^ and motif fragmentation happened in the first layer of deeper DCNN^14^. But the kernel size in the first layer has to be large enough to learn a single complete motif^5^. However, deeper DCNNs show better performance in genomics^15,16^. Hence, performance and interpretation seem to be a trade-off determined by the DCNN architecture to a large extent.

Here, we proposed NeuronMotif to decipher transcriptional cis-regulatory grammar from DCNN (Fig.1 d,e). This algorithm considers the sequences recognized by an Artificial Neuron (AN) as a mixture model depending on latent variables. From the output of AN to the input, it automatically backward discovers the latent variables reflecting the neural network structure to decouple the AN mixture model for extracting motif grammar. We applied NeuronMotif on several existing shallow DCNNs (DeepSEA^4^ and Basset^5^). A large portion of uncovered motifs and syntaxes of their combinations are supported by literature or ATAC-seq profile, which outperforms existing state-of-art methods. The results of NeuronMotif reveal the origin of adversarial noise in the model, which can be used to guide the design of DCNN architectures to suppress noise. With the help of NeuronMotif, we further built and interpreted 10-convolutional-layer deeper DCNNs with the help of NeuronMotif.

## Results

### The NeuronMotif algorithm for uncovering motif and decoupling motif mixture from DCNN model

TFs are proteins that can recognize and bind to specific DNA sequences. The perferred sequences bound by a given TF are usually summarized as a motif. Motif is a model typically refers to Position Weight Matrix (PWM), which can be converted from Position Probability Matrix (PPM)^17^. At each base position in PPM, the four scores represent the probability of the four bases that occur at the relative position of TFBS. The probability can be estimated by collecting the DNA sequences binding with TFBS through experiments such as Systematic Evolution of Ligands by Exponential Enrichment (SELEX)^18^ (Fig. 2a). This process is similar to sampling sequences (x) of TFBSs with *N-* bases length from an 4 × N random variable matrix ***X*** ~ *p*_**PPM**_4x*N*__(***x***) to estimate PPM 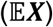 by element-wise average 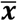 (see Methods). Here, ***x*** is the 4 × *N* one-hot code of the sequence, and each column of ***X*** is an different independent categorial distribution. The sampling process in the experiment reflects TF binding affinities to sequences. The sequences with stronger affinities may occur at higher frequency.

**Fig. 2.**
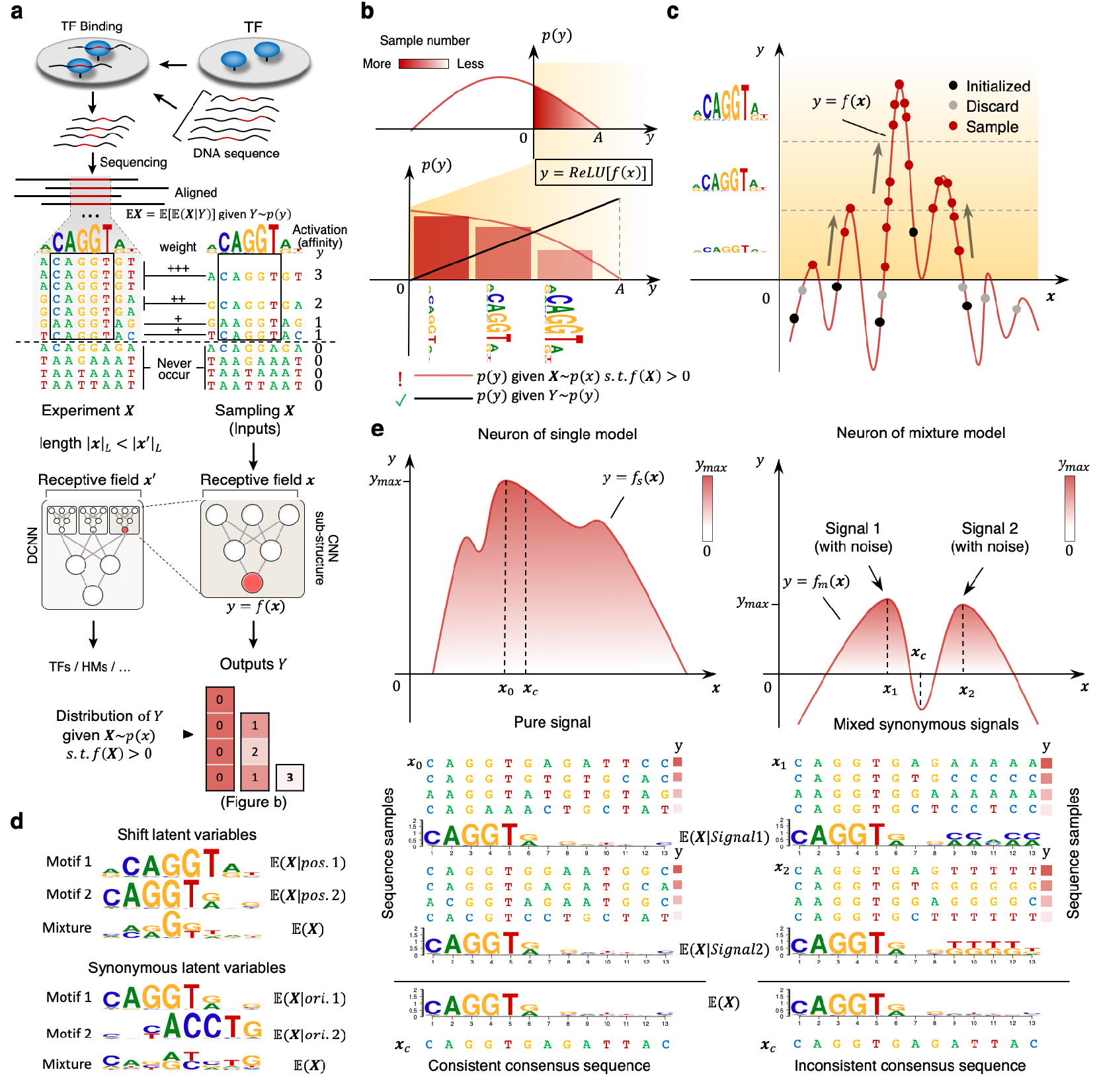
Details of NeuronMotif. **a**, In experiment like SELEX, the sequences (z) bound by TF are filtered and aligned for motif estimation 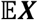. To simulate this process, enumerating the valid sequence (x) for estimating the motif for a neuron is not correct (given distribution of X). The frequency/weight of sequences should be proportional to affinity level (given distribution of ***Y*** in **b**). Within the whole DCNN structure, DCNN substructure of the neuron in red is equivalent to function *y* = *f*(***x***). The abbreviation *s.t.* means subject to. **b**, Distribution of neuron activation values (*y*) in **a**. The sequence collection with a higher activation level contains more information in the sequence logo. **c**, The sequences are sampled during the optimization process of seed sequence. **d**, Two types of latent variables lead to motif mixture in neuron model. The shifted motifs can be decoupled under the control of shift latent variable that determine the position 1 and 2. The synonymous latent variable determine the different replaceable motif with similar function at the same position. The example is the original motif 1 and its reverse complementary motif 2. For some TFs, function is not sensitive to orientation. **e**, Comparing the neuron with (left column) and without (right column) synonymous mixture motif. Under the controlling of synonymous latent variables *S* = *S*1,*S*2, the sequences and corresponding motifs are similar in single model but different in mixture model. The sequences with max activation value in two model are ***x*_1_,*x*_2_,*x*_3_**(*f_s_*(***x*_1_**) > *f_m_*(***x*_2_**) ≈ *f_m_*(***x*_3_**)). Both of the models share consensus sequence (x_c_).

Inspired by SELEX screening TF-preferred sequences, we attempted to imitate this process by sampling AN-preferred sequences to study AN. The sub-structure of an AN processes the sequence input (x) with a non-linear function *y* = *f*(***x***) and then outputs an activation value (*y*) (Fig. 2a and Fig. S4a). This process is quite similar to SELEX screening sequences because the sequences (***x***) with higher activation *y* are preferred by the AN for affecting downstream ANs and the final prediction result, which reflects sequence affinity. Hence, the input random variable matrix ***X*** ~ *p*_PPM_4×*N*__(***x***) depends on the output random variable *Y* ~ *p*(*y*) through *Y* = *f*(***X***). To obtain PPM reflecting binding affinity rather than binding probability, we adopt a linear function as *p*(*y*) of the distribution (Fig. 2b, see Methods for detail explanation). In other words, sampling weight or frequency of each unique sequence (***x***) should be positive proportional to its activation value (*y*) (Fig. 2a and 2b). It can be implemented by sampling ***X*** at the same level of *y* to estimate 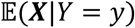 (bottom of Fig. 2b) and then taking the weighted average of them 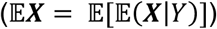 to estimate PPM (Fig.2b, see Methods for details). This method precedes previous studies in representing the TF binding affinity to DNA sequences. Adapted Back Propagation (BP) methods like Saliency Map and DeepLIFT do not model the sequence preference with ***X***. The importance score (e.g. *∂y*/∂***x***) of these methods do not directly reflect PPM or PWM (Fig. 1c). While adapted max activation methods like the methods developed by Kelley et al. and Alipanahi et al. for interpreting Basset model^5^ and DeepBind model^8^ try to follow the PPM model but they estimate 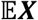 by 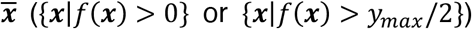 without depending on the level of *Y*, thus could hardly reflect the activation perference.(Fig. 1b). Here, we assumed that TFBS are located at the same relative position in the input sequences without shifting, and then we can define motif or PPM for the sequences recognized by an AN as 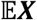 given distribution of *Y.* We called it AN motif or PPM of AN.

However, we found that due to the max-pooling operation in DCNN, TFBSs may be located at different relative positions in the input sequences to activate the AN. In the max-pooling layer, the key input feature maps reflecting the shifting diversity of TFBSs will be unified into similar output feature maps (Fig. 1d, 1e and S4b). The downstream key ANs including the output AN will share similar activation values (*y*) for different sequences with shifting TFBSs. Hence, the motif sequences of TFBS recognized by an AN can be regarded as a latent variable mixture model. To decouple motif mixtures, we have to find some shift latent variables that reflecting different positions of the motif sequence (the top part of Fig. 2d). Only by controlling these latent variables can we obtain the consistent real sequence motifs 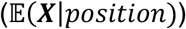. This key issue is neglected by all existing methods (Fig. 1b, Fig. 1c and Fig. S1).

We further found that TFBSs in the sequences may not share the same pattern. It indicates that we can find more than one motif by stacking TFBSs with grouped consistent pattern respectively. One of the cases is the reverse complementary sequences (bottom part of Fig. 2d). The mixing of these sequences can be controlled by another important type of latent variables in the mixture model named as synonymous latent variables, and we called the decoupled motifs as the synonymous motifs. The synonymous motifs represented by an AN should satisfy: (1) they are not shifted motifs; (2) all or part of input variables ***X*** are conditionally independent under controlling synonymous latent variables; (3) the sampled sequences grouped by synonymous motifs should share similar maximun activation values so that they are all preferred by the AN. If these conditionally independent positions affect little on activation values, then the AN can be regarded as a single model (SM, *y* = *f_s_* (***x***), the left column of Fig. 2e). Otherwise, the motif sequences recognized by the AN is a mixture model (MM, *y* = *f_m_*(***x***), the right column of Fig. 2e). Both SM and MM share similar motif (the bottom part of Fig. 2e), but the sequences of the maximum activation value (SM:*x*_0_; MM: *x*_1_,*x*_2_) and the consensus sequences (***x***_*c*_) show their difference. Different from *f_s_*(***x***_*c*_) ≈ *f_s_*(***x***_0_) in the SM, *f_m_*(***x***_*c*_) in MM usually strongly deviates from *f_m_*(***x***_1_,*f_m_*(***x***_2_), and could even be negative (the top-right part of Fig. 2e). This is because ***x***_*c*_ may not match any conditionally dependent motifs embedded in the AN (the bottom-right of Fig. 2e). Thus, bases flipping at the conditionally dependent positions of the sequence is a kind of adversarial noise^19^ discussed in CV that can dramatically change the AN activation level or even destroy prediction result. In addition, we also proved that severe mixing of synonymous motifs in the AN correlates to the lower maximum activation and weights of the AN (see Methods). The results above suggested that the mixture of synonymous motifs seems to be noise rather than motif signals due to its vulnerable characteristics. Hence, for a well-trained model with weak noises, we only need to decouple the signal of each AN depending on the max-pooling structure.

One of the most widely-used types of DCNN models is composed of general convolution layers and max-pooling layers. We took this type of DCNN as an instance, and developed the NeuronMotif algorithm to uncover the motif combinatorial grammar from DCNN. First, we designed a sampling algorithm adapted from genetic algorithm to optimize seed samples and recorded the intermediate valid sequences as the sampling result (Fig 2c, see Methods for details). Second, we used K-means (K is pooling size) to decouple the mixture signal from different sequences by clustering the shifting similar sub-patterns in the input feature map of the max pooling layer to split the sequences set (Fig 1d,e and Fig. S4b). The decoupling process can be performed backward and recursively from the deepest layer to the first layer. Third, the algorithm can annotate an AN with motifs by estimating 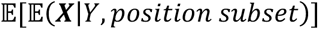 from each subset samples clustered by K-means (Fig 1e, see Methods for detail).

The steps above can only decouple the mixture of a single hard AN motif with shifting diversity. The hard motif refers to the motif or motifs combination with a fixed gap, which characterizes homodimer, heterodimer or multimer TFs that can be considered as a stable molecular cluster binding to DNA. However, a large portion of TFs cobinding are gapped by flexible intervals. Their sequence pattern is the soft motif that composed of more than one hard motif, and the space between any two adjacent hard motifs is in a certain range.To decouple the hard motifs in a soft motif represented by AN, users should run the decoupling algorithm in NeuronMotif (the second step) iterately for several times based on the number of hard motifs (Fig 1e, see Methods for details).

### NeuronMotif successfully decouple the motif mixture

To evaluate the performance of NeuronMotif on decoupling the motif mixture signal, we applied NeuronMotif to annotate two well-known models, DeepSEA^4^ and Basset^5^, both of which are DNA-sequence based DCNN models with 3 general convolutional layers for genome function annotation. Basset annotates open chromatin region trained by DNase-seq data. In addition to chromatin accessibility, DeepSEA also annotates TFBSs and HMs trained by ChIP-seq data. NeuronMotif successfully decoupled the shifted mixture motifs from layer 2 (L2) and layer 3 (L3) of the both models (see Supplementary Information for all results). In the Basset model, the first- and second-layer pooling size are 3 and 4, so the numbers of shifted signals are 3 and 3 × 4 = 12 for L2 and L3 ANs, respectively (Fig. 3a-c). In the DeepSEA model, the first- and second-layer pooling size are both 4, so the numbers of shifted signals are 4 and 4 × 4 = 16 for L2 and L3 ANs, respectively. In Fig 3, all adjacent AN motifs are shifted with 1bp and highly consistent. However, the state-of-the-art methods, such as Kelley et al^5^., Alipanahi et al^8^ and Saliency Map^20^, cannot deal with the mixture signal which leads to the much lower information content and very noisy signals.

**Fig. 3.**
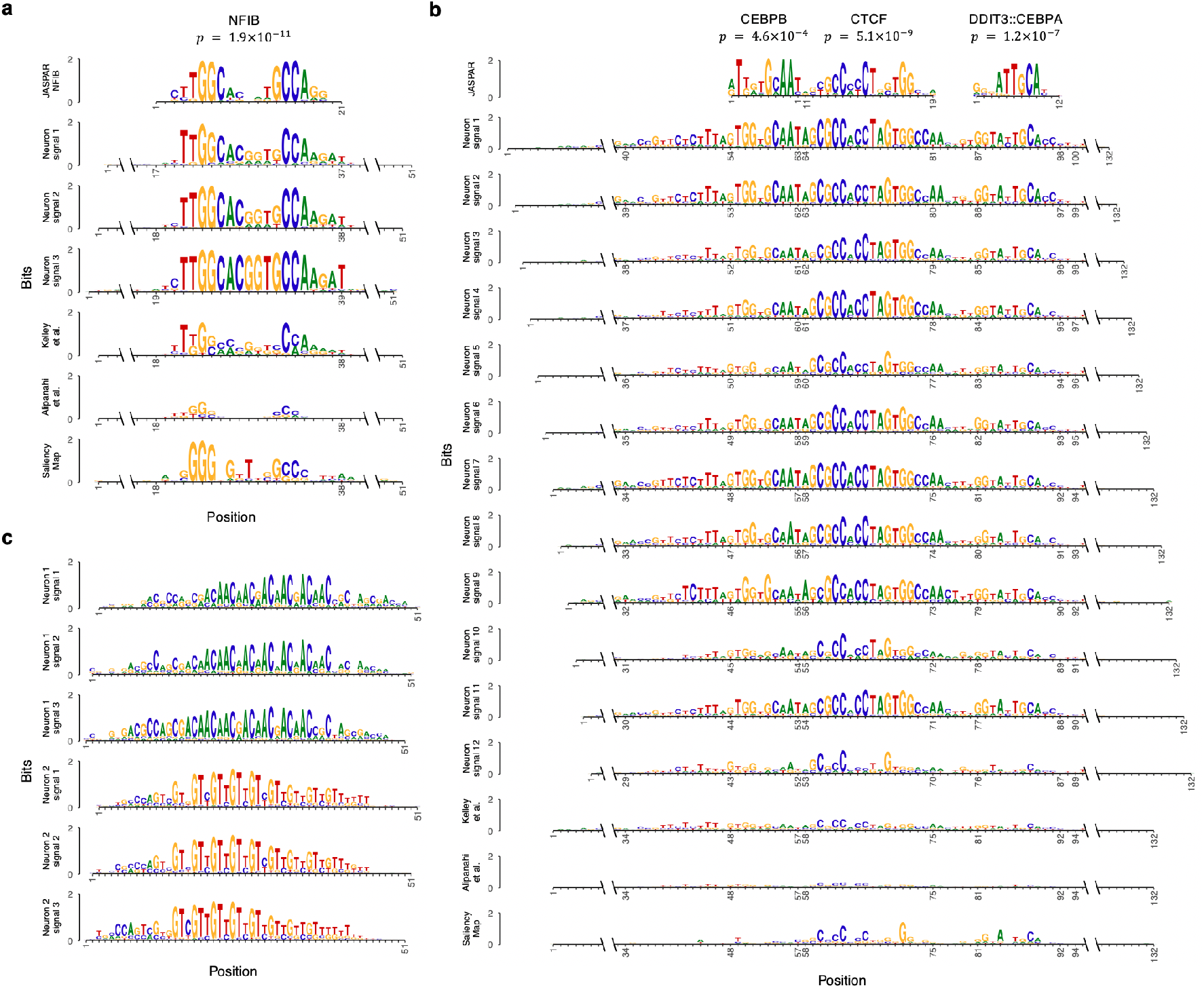
Use NeuronMotif to annotate Basset model. **a**, Four motifs of a second-layer neuron decoupled by NeuronMotif (row 2-4). The decouple motifs with the same size (the receptive field size of neuron in the second layer is 51bp) are aligned with 1bp offsets. They are matched by JASPAR motif NFIB using Tomtom (row 1). The interpretation results using methods of Kelley et al., Alipanahi et al. and Saliency Map are shown in row row 5-7. **b**, Motifs of a third-layer neuron decoupled by NeuronMotif (row 2-13). The decoupled 12 motifs with the same size (the receptive field size of neurons in the third layer is 132 bp) are aligned with 1bp offsets. They are matched by JASPAR motif CEBPB, CTCF and DDIT3::CEBPA using Tomtom (row 1). The interpretation results using methods of Kelley et al., Alipanahi et al. and Saliency Map are shown in row row 14-16. **c**, The 2 neurons in the second layer learn the reverse complementary motifs. They represent the motifs of AAC triplet repeats (row 1-3) and GTT triplet repeats (row 4-6) respectively, which were decoupled by NeuronMotif.

The NeuronMotif-annotated results of Basset and DeepSEA models showed that ANs extract various kinds of motifs. Here, we took Basset as an instance. Some ANs extract important TF motifs correlated with Basset’s prediction targets (DNase I sensitivity). Many motifs of ANs can be matched with the known motifs in the JASPAR^21^ database (Fig. 3a and 3b). Some matched TFs, like NFI and cobinding TFs CTCF-CEBP, are highly correlated with chromatin openness^22,23^. In comparison, the interpretation result of existing methods can hardly be matched with any known motifs in JASPAR. Statistically, NeuronMotif found more motifs and more accurate motifs from JASPAR database (Fig 5a and 5c). Besides, some important functional sequence features and their reverse complement can also be identified from motifs of AN. One of the typical examples is the repeats of AAC triplets feature extracted by the Basset model (Fig. 3c). It has been reported that repeated triplets AAC is enriched in intron^24^. As the intron regions are usually open for gene transcription, it is reasonable that the Basset model extract this feature.

### DCNN diagnosis and architecture design guidance

From the NeuronMotif result of DeepSEA and Basset models, we found that the outputs of some ANs were always zero no matter how we changed the input sequences. We called them dead ANs (Fig 4a). The dead ANs are redundant because they cannot affect the downstream network. During the sampling process of annotating DeepSEA model via NeuronMotif, we found the sampling algorithm cannot sample even one sequence that can activate some ANs in L2 and L3. For example, A total of 150 and 120 ANs in L2 and L3 are dead ANs in the DeepSEA model.

**Fig. 4.**
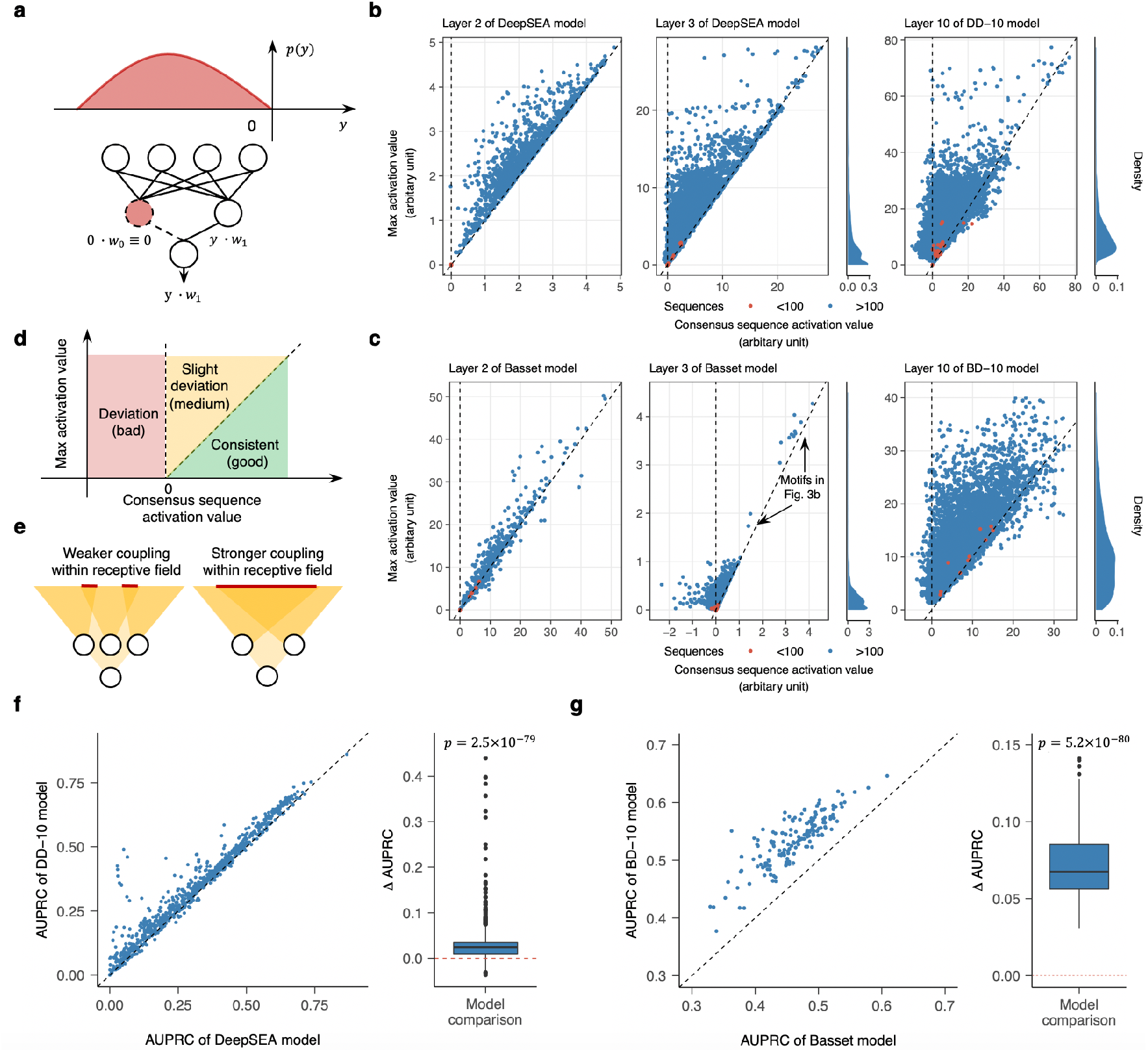
NeuronMotif diagnose model defects and guide DCNN architecture design for better performance. **a**, Dead kernel definition. Dead neuron (pink) activation value distribution is negative. It will be filtered by ReLU activation function. The output of dead neuron is zero. The downstream neuron output does not depend on this neuron. **b**,**c**, Diagnosis of motif mixture in the decoupled motifs from Basset, BD-10, DeepSEA and DD10 model. Each point is a decoupled motif generate by a sample set of sequence. The points of motifs generated by the sample set with less than 100 sequences are marked by red color. Otherwise, they are marked by blue color. The distribution of the max activation value is used to show if the relative max activation values of most of the motifs are too low. **b**, Diagnosis of models trained by DeepSEA dataset **c**, Diagnosis of models trained by Basset dataset. Only the max activation value of the decoupled motifs in Fig. 3b are significantly higher than the decoupled motifs of other neurons in layer 3 of Basset-3 model. **d**, The meaning of each region in the sub-plots of **b**,**c**. **e**, Schematic for receptive field coupling of previous layer neuron in the neuron sub-structure. **f,g**, Use AUPRC as an indicator to compare the prediction performance of models. For each model pair, one-sided t-test of Δ*AUPRC* = *AUPRC_y-axis_* – *AUPRC_x-axis_* is used to access model performance difference level. **f**, Comparison between DeepSEA and DD-10 models. **g**, Comparison between Basset and BD-10 models.

Another problem is that some ANs may recognize synonymous motifs. We diagnosed this problem with two indicators of motifs based on the phenomena that an AN may represent synonymous motifs. One indicator is the activation value of motif consensus sequence and the other is the maximun activation value of the sampled sequences for motif estimation. We found that if the two indicators severely deviate from each other, or the max activation value is close to zero, then the corresponding AN may suffer from the synonymous motifs problem. The two indicators of Basset L2 ANs are almost consistent (the first column of Fig. 4c). However, in L3, activation values of many decoupled motifs’ consensus sequences are negative and severely deviated from the maximum activation value (the second column of Fig. 4c). Most problematic motifs are mainly caused by stacking sequences of different synonymous motifs with the maximum activation value closed to zero (see Methods for details and Supplementary Information for case).

To overcome these problems, the DCNN architecture should be optimized to avoid the mixed signals of synonymous motifs. Both of DeepSEA and Basset use large convolutional kernel size (> 8) and large pooling size (3 or 4) in each layer. For an AN with the certain receptive field, implementing its sub-structure with larger kernel size and pooling size tend to cause weaker coupling among sub-structures of the previous layer ANs (Fig. 4e). When using the same training set and optimization method for training the model, we found that the less coupling among the ANs, the more sensitive to noise generated by synonymous motifs (see Methods and Fig. S6). DeepSEA adopted strong regularization methods to successfully suppress learning these the noises (Fig. 4b) but with the cost of producing dead ANs. In the field of CV, building deeper networks with smaller kernels and pooling structures has been found to be a more robust strategy with better performance^25^. Thus, we built 10-convolution-layer new models and trained them on the Basset Dataset (BD-10) and the DeepSEA Dataset (DD-10) respectively. The synonymous motif problem was significant suppressed in BD-10 and DD-10 (the third column of Fig. 4b and Fig. 4c), and few dead kernels were found. Furthermore, Both BD-10 and DD-10 show much better prediction performance (Fig. 4f and Fig.4g) than the original model. These results demonstrate how NeuronMotif can be used to help diagnose DCNN and guide architecture design.

### Accuracy and completeness of motif discovery in different layers of DCNN

To study which layer is better for motif discovery in a DCNN model, we used NeuronMotif to interpreter the shallow convolutional layers with receptive field around 19bp and the deepest convolutional layers in three models with 3, 5 and 10 convolutional layers (Basset, BD-5 and BD-10) trained by the same data of Basset paper. To measure the interpretation performance, we matched the decoupled motifs to the motifs in the JASPAR^21^ database using Tomtom^26^. For each AN matched to known motifs (*q*-value < 0.1), we selected the best matched motif in JASPAR and took similarity measurement between the found motif and the JASPAR motif (*q*-value) as the performance of the AN. As the numbers of ANs are different in each layer, we only selected the q-values of top 100 ANs for further analysis. Given a DCNN model, we found that the motifs discovered from the deepest convolutional layer outperform the shallow layers with around 19bp receptive field (each column in Fig 5a). We further compared the layers with similar receptive field (the first row in Fig 5a), and the deepest convolutional layers (the second row in Fig 5a) among different models. The deeper models (BD-5 and BD-10) outperform the Basset model by discovering much more known motifs (Fig. 5b) and the motif is matched better to the JASPAR database (each row in Fig 5a). Based on the comparison results, we recommand to use deep AN for motif discovery and representation. For example, we built a motif dictionary from layer 10 (L10) in BD-10 model. This dictionary contains 9056 motifs. Among them, 7974 motifs are matched to at least one of 399 JASPAR motifs by Tomtom (q-value < 0.01, one of the best-discovered motifs matched to each JASPAR motif is shown in Fig 5d) and remaining 1082 motifs are novel motifs.

**Fig. 5.**
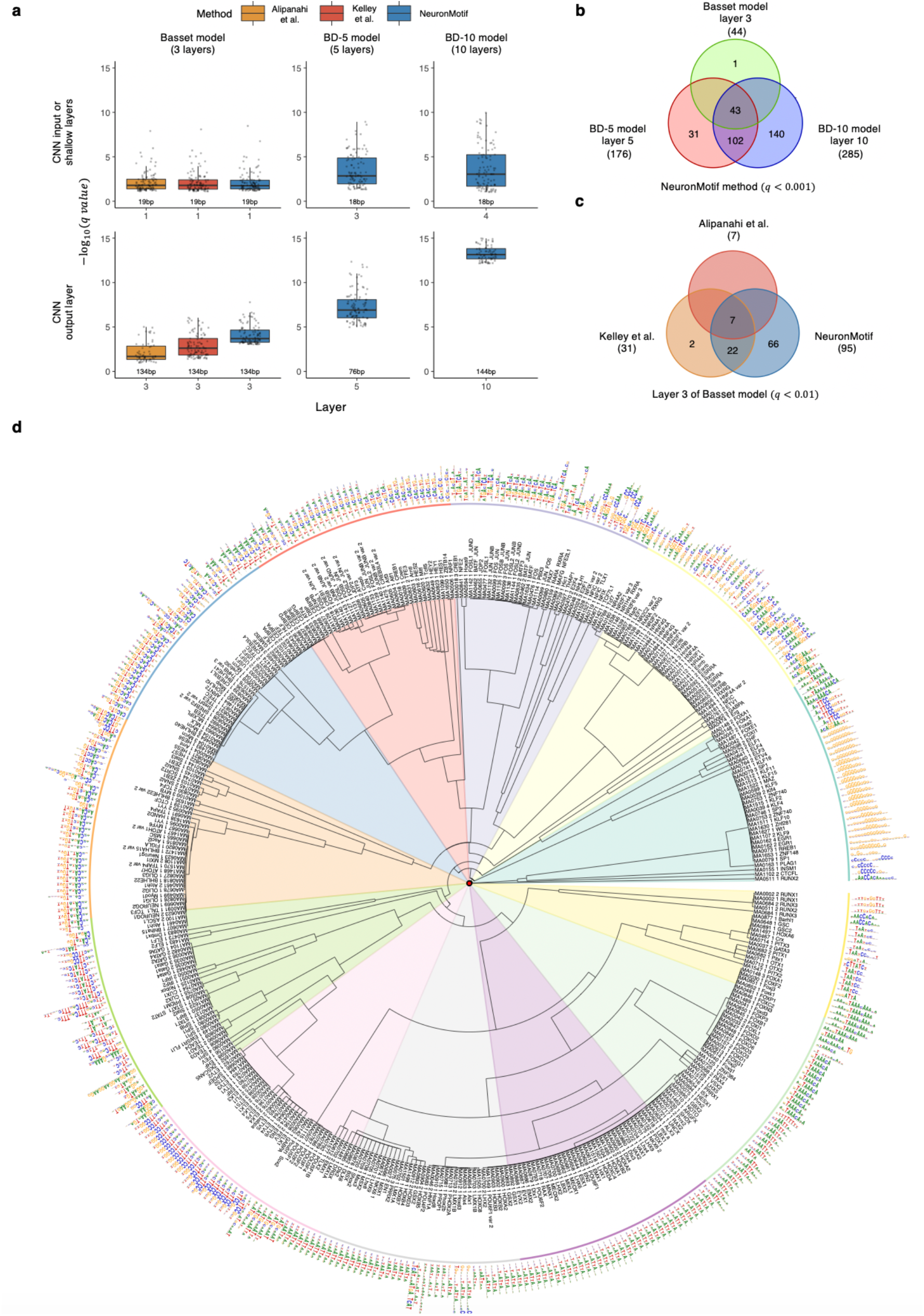
Motif discovery performance for different layers in different models. **a**, Accuracy analysis for discovered motifs of different models. Three columns of box plots describe similarity between neuron motifs and JASPAR motifs in Basset, BD-5 and BD-10 model respectively. For each selected layer in the model, -log10(q-values) distribution of the top 100 JASPAR motifs matched neuron (q-value < 0.1) are shown with box and jittering points. The color of the box means the applied interpretation method. In the first row, it shows the result of input (first) layer of Basset model and shallow layers in BD-5 and BD-10 model with similar receptive field sizes (18 bp) of the Basset input layer neuron (19 bp). In the second row, it shows the convolutional output layer, the convolutional layer in the front of dense layer, result of the three models. **b**, The number of motifs discovered (q-value < 0.001) from the neuron in convolutional output layer of Basset, BD-5 and BD-10 model. **c**, The number of motif discovered (q-value < 0.01) from the neuron in layer 3 of Basset model using different interpretation methods including Kelley et al., Alipanahi et al. and NeuronMotif. **d**, Discovered motifs from the neuron of top convolutional layer in BD-10 model (q-value < 0.01). These motifs can be matched to JASPAR database. Only the one with smallest q-value for each JASPAR motif is shown.

### NeuronMotif successfully uncover motif grammar

In previous works^21,27^, motif combination grammar is usually represented by the hard motif. They depict soft motif by enumerating different intervals among the component of hard motif (Fig 5a and 5b). In comparison, DCNN structure is more powerful to describe these soft motifs when the receptive field is long enough. Here, we take L10 ANs in DD10 as examples to study the AN soft motif. We assume the L10 ANs representing no more than two hard motifs, so we run decoupling algorithm twice in NeuronMotif and a total of 256 AN motifs are generated (Fig. 6a and 6b, see Methods for details). These AN motifs enumerating the combination of hard motifs with various sizes of gap. From these AN motifs, we can slice the shared hard motif to build motif dictionary. Based on the dictionary and all AN motifs, we can summarize the interval range between arbitray two adjacent hard motifs and build the syntax tree (Fig. 6a and 6b). Some of the soft motif can be supported by literature. For example, an AN in DeepSEA represents the soft CTCF homodimer with around 58 bp interval that play important roles in the transcriptional process of cancer and germ cells development^28^ (Fig. 6a). We also found that DDIT3::CEBPA can co-bind with CTCF, which is not reported in previous literatures. Interestingly, CTCF-DDIT3::CEBPA is shown to be an conservative hard trimeric motif that also occurs in the Basset model (Fig. 3b), which show the reliability of this discovery. We further used the ATAC-seq data footprinting to validate the discovered AN motif grammars. ATAC-seq uses Tn5 transposes to cut DNA into fragments. If there are some TFs or other molecules binding to DNA, the cutting frequency will be affected. For each AN, we aligned corresponding Tn5 transposes cutting frequency of top 3000 sequences (144bp) with max AN activation values in the test dataset. We extended the footprinting region to 1000bp in total. Most ANs have their own footprintings generated by ATAC-seq data from five cell types or tissue (Fig. 6c and Fig. 6d, see Supplementary Information for other ANs). Soft CTCF-DDIT3::CEBPA homodimer footprintings from five cell types or tissue share the pattern of three peaks and two valleys (Fig. 6c) but soft NFI homodimer footprinting signals are only significant in prostate tissue and LNCaP cell lines which support the notion that NFI family can regulate prostate-specific gene expression^29^ (Fig. 6d). The results indicated that some motif grammars of multimers are cell-type specific. To further confirm the footprinting caused by the specific TF binding, we calculated the distribution of motif matched positions for both CTCF–DDIT3::CEBPA and NFIC motif (Fig. 6e and Fig. 6f). The peaks of motif-matched positions are consistent with the footprinting valley. All the results suggested that NeuronMotif provides a novel way to discover the soft multimer motif grammar on the genome and to better depict multimeric TF motifs.

**Fig. 6.**
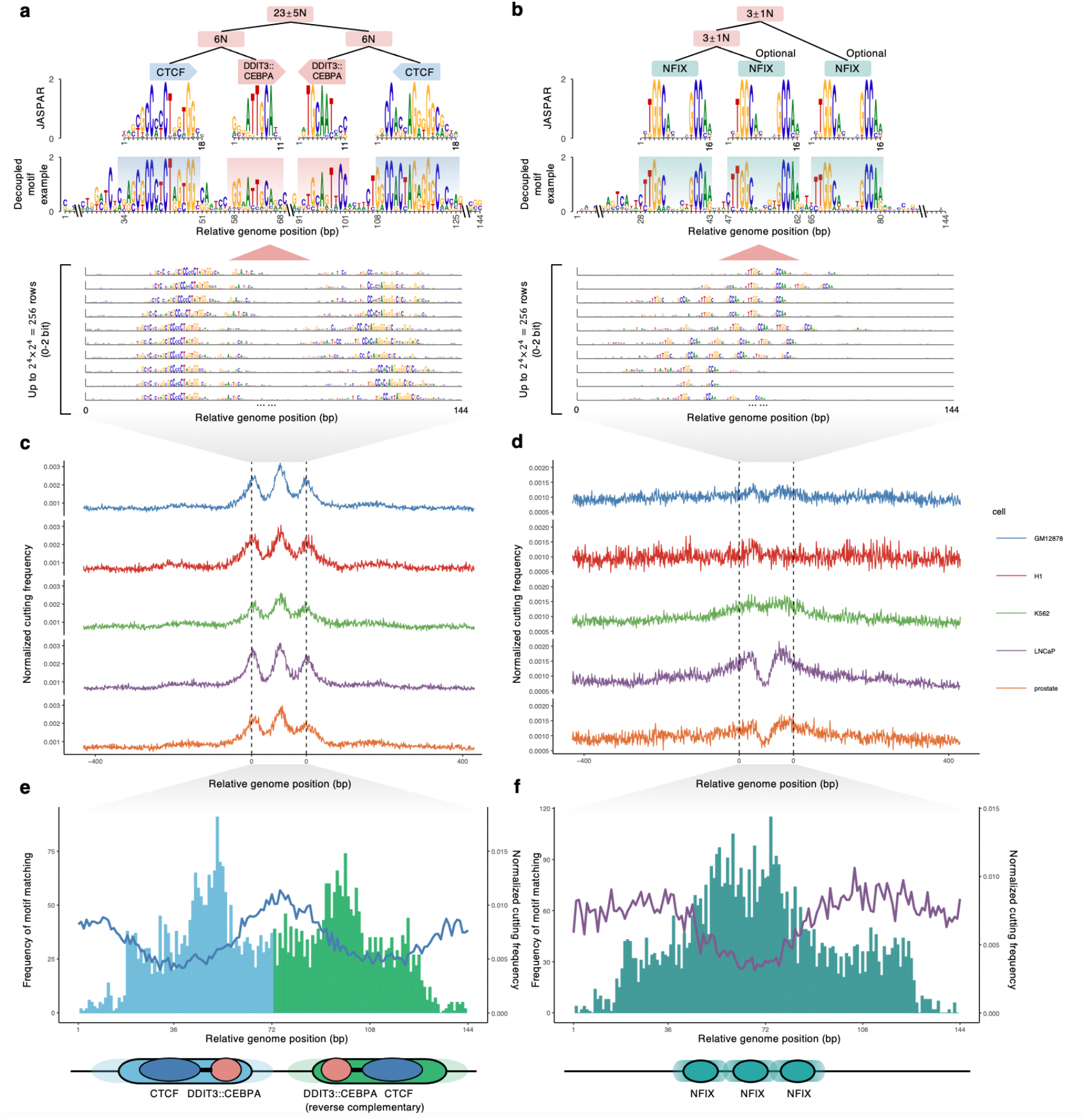
Verification of neuron motif. **a**, Motif syntax represented by a neuron of top convolutional layer (layer 10) in DD-10 model applied on DeepSEA data. The neuron motifs (144 bp) are matched to TF CTCF and DDIT3::CEBPA. CTCF-DDIT3::CEBPA is a hard hetero-trimer. The distance between two CTCF-DDIT3::CEBPA trimer is flexible. **b**, Motif syntax represented by another neuron of top convolutional layer (layer 10) in DD-10 model applied on DeepSEA data. The neuron motifs (144 bp) are matched to TF NFIX. **c**, Five different cell types’ ATAC-seq data footprinting (500 bp upstream and downstream from the motif matched midpoint is shown) of the motifs in a. Cut-site counts of each position are normalized by total cut-site counts within 1000 bp window. **d**, Similar to **c**, the footprinting of the neuron motif in **b**. **e**, CTCF-DDIT3::CEBPA motif matched count for each relative motif midpoint position. Soft homodimer of CTCF-DDIT3::CEBPA heterotrimer relations are shown at the bottom.. **f**, NIFX motif matched count for each relative motif midpoint position. Soft homotrimer of NIFX relation is shown at the bottom.

## Discussion

In summary, we presented NeuronMotif as an effective algorithm to reveal the cis-regulatory motif grammar learned by DCNN model that use DNA sequence to annotate genome function. We proposed the statistical form of AN motif representation and the latent variable mixture model to understand each convolutional neuron. Take maxpooling-convolutional structure as an instance, we uncovered the signal mixing mechanism including shifting latent variable and synonymous latent variable. The NeuronMotif used a K-means-based algorithm to decouple the latent variable mixture, and a sampling strategy adapted from genetic algorithm for motif estimation. We eveluated NeuronMotif interpretation performance on DeepSEA, Basset and some inhouse deeper models. Many uncovered motif conbinatotial grammars are supported by literature and ATAC-seq data. Finally, we showed that NeuronMotif result can be used for model diagnoses and to guide model structure design for better prediction performance and motif extraction.

Except for interpretating cis-regulatory motif grammar from DCNN, the application of NeuronMotif may be extend to many other problems. DNA sequence is a special onedimensional discrete data with four elements. It is possible to apply NeruonMotif to the DCNN for amino acid sequence of protein or other continuous sequence like different kinds of sequencing profile. There are still some issues that should be addressed to further expand the application of NeuronMotif. For instance, NeuronMotif only focuses on max-pooling-CNN structure. Many new DCNN structures such as ResNet and DenseNet are put forward in recent years. As these structures show better performance in CV, it is valuable to adapt the NeuronMotif method for these more general and complex DCNN structures in genomics studies.

In the future, we envision that DCNN model interpreted by NeuronMotif will advance our ability to discover and summarize the complicated regulatory rule, model transcriptional cis-regulatory process and understand DCNN blackbox itself.

## Acknowledgements

We thank Z. Duren and H. Fang for valuable suggestions on motif discovery and relative biological issues. This work was supported by the National Natural Science Foundation of China (No. 62050152, 61721003), and the National Key R&D Program of China (No. 2020YFA0906900)

## Competing interests

Tsinghua University has a patent pending for NeuronMotif.

## Author contributions

Z.W., W.H.W. and X.W. conceived the main idea of the study. Z.W. completed the theorem proof and formula derivation. K.H. repeated and checked the proof and inference. Z.W. developed the algorithm, trained DCNN model, designed experiments and implemented all the experiments. R.J. provided and maintained the computing cluster. W.H.W. and X.W. designed some experiments and supervised the study. All authors wrote and revised the manuscript.

## Methods

### Statistical definition and estimation of PPM represented by a convolutional neuron

An AN has its own sub-structure in the DCNN model (Fig. 2a and S4a). The substructure includes an input ***x*** and an output *y*. The relation between ***x*** and *y* is defined by a non-linear function *y* = *f*(***x***) which depends on the sub-structure of the AN, because all upstream ANs in sub-structure can affect the characteristic of AN. The input of each AN is the output of AN in the previous layer. The sub-structure also determine the receptive field size *N* (the length of ***x***).

For each valid DNA input sequence (***x***,s.t. *f*(***x***) >0), ***x*** is a 4×*N* matrix of one-hot code. It can be sampled from a random variable matrix ***X*** ~ *p*(***x***). *X* contains 4×*N* random variables ***X***_*b,j*_ (*b* = A,C,G,T; *j* = 1,2, …,*N*). Each column can be modeled as an independent multinomial distribution (***X***._*j*_~Multi(l,***π._j_***)), where the 4×*N* probability matrix ***π*** is the PPM that characterizes the preference of the nucleotide bases for the sequence motif. Based on the nature of multinomial distribution, the parameter 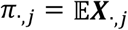 so PPM can be estimated through sampling ***X.***_,*j*_ and calculating the element-wise average 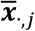. As the unknown distribution *p*(***x***) is to be estimated, we cannot sample ***X*** directly. We know that ***X*** is not a free random variable, but depends on the free output random variable *Y*~*p*(*y*) through *Y* = *f*(***X***). Based on the identity equation 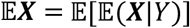, we can first sample ***X***|*Y* = *y* to estimate 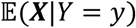, which represents the PPM for a specific activation value or affinity (*y*). Given an arbitrary distribution *p*(*y*), we can obtain the PPM by taking a weighted average of these PPMs with different affinities.

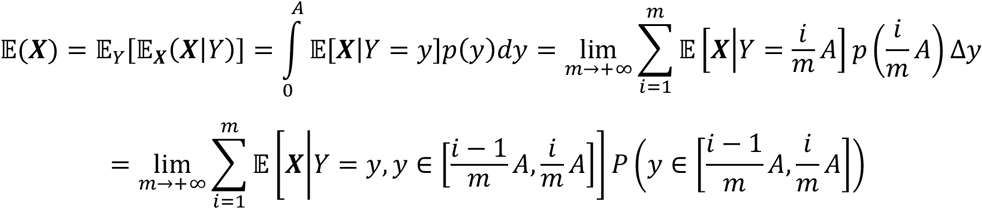

Numeric estimation of PPM for an AN needs enough valid sequence (***x***) samples. In this work, we set ReLU(*f*(***x***)) = max{*f*(***x***), 0} as the activation function of each convolutional neuron. Therefore, the valid sequence dataset is ***X***^+^ = {***x***|*f*(***x***) > 0 ∧ |***x***|_*L*_ = *N*}. Here, *f*(***x***) > 0 constrains the activation value of a valid sequence to be positive so that it can activate the AN, and |*x*|_*L*_ = *N* constrains that the length ***x*** must match the AN receptive field size *N.* For convenient, we rewrote the sequence dataset as 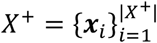 and corresponding activation value set as 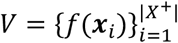. The max activation value is *A* = max(*V*). The probability of TF binding to a DNA sequence depends on the binding affinity ^30^. To sample the sequences reflecting their affinity levels (*y*), the sequence with high affinity should be sampled in higher frequency. Here, for the ease of calculation, we set the probability density function of *Y* as a linear function 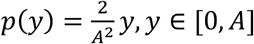 (Fig. 2b).

In practical, we split interval [0,*A*] into *m* (*m* = 20) bins to merge sequences with similar activation values into PPMs (Fig. 2b). In this way, we can get the average of PPMs weighted by activation values. For each bin *i* (*i* = 1,2, …,*m*), the sequence index set is 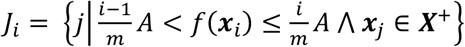. Sequences in bin *i* share similar activation values. Thus, their average activation value and PPM can be calculated by

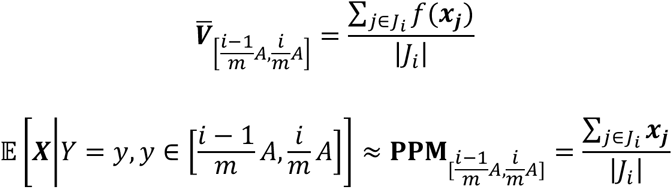

where |*J_i_*| is the number of sequences in sequence index set *i*. The probability or weight for each bin can be estimated by

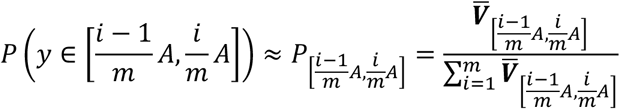

Finally, **PPM**_4x*N*_ of the AN can be estimated by the average of PPMs weighted by the activation value.

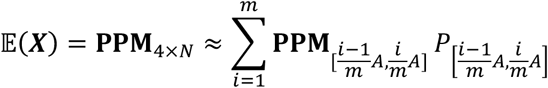

The estimation above assumes that relative position of TFBS in the input sequence are the same and all of them share the same motif. In other words, it only works for SM neurons. However, the assumption was not suitable for most ANs especially ANs in deeper layer, where the estimation result is a mixture of different motifs. The random variable matrix ***X*** can be considered as a MM. Hence, we first needed to find the latent variables that can split the dataset ***X^+^*** into subsets 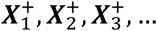, each of which can be consider as an SM. The estimation should be applied on each subset respectively. It will generate several motifs **PPM**_1_,**PPM**_2_,**PPM**_3_,… which are controlled by different conditions of the latent variables.

### Discovering latent variables in a mixture model of neuron

The activation value of the AN is the only indicator to show the matching level of a sequence. Sequences with high activation values of an AN may be composed of completely different key TFBSs at various relative positions due to the powerful representation ability of the neural network. This characteristic shows that sequences recognized by an AN can be considered as a latent variable mixture model. The sequences matched by different sub-models in the MM are available to activate the AN at the same level. Hence, the activation value of the AN is the unified or mixed signal that cannot distinguish the sequences with different TFBSs. To find the mechanism of mixing process for an AN, we can investigate the activation values of all upstream ANs (feature maps) that reflects TFBS diversity in the sequences, which define the latent variables. Controlling these latent variables, the sampled sequences share the same pattern (hard motif). The sequences are only matched by one sub-model in the MM. In this way, different sampled sequences shared the similar TFBSs that are located at the same relative position. Only obtain these sampled sequences can we estimate the AN motif.

In practical, when analyzing the feature maps for sampled sequences, we used K-means to cluster the feature maps of the convolutional layer and found shifted signals among each cluster. However, these clusters are not able to be rebuilt by the feature map of the downstream max-pooling layer. So, the max-pooling operation unify the shifted signals of various sequence, which removes the difference among clusters. In other words, AN just tries to detect if TFBS exist in sequence, the position of TFBS in the sequences is not important to final AN output. Subsequently, we found the best cluster number K (the maximum shifting offset) is the same as the max-pooling size. Each offset within K defines a shifting latent variable.

### The side-effect for the ANs representing different synonymous motifs

Following the definition of synonymous motifs for an AN, if an AN (*y* = *f*(***x***)) represents the mixture of two synonymous motifs, let ***x***_1_, ***x***_2_ be the vectors of flattened one hot code of the maximum activation sequences for the two motifs respectively, then they should satisfy *f*(***x***_1_) ≈ *f*(***x***_2_) i.e. *f*(***x***_1_) - *f*(***x***_2_) → 0. First, we studied the AN in the first layer (*y* = *f*(***x***) = ***kx***^*T*^ + *b*). The activation values of ***x***_1_,***x***_2_ are

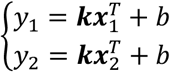

Where ***k*** is the weight of the AN, and *b* is the bias or inceptor. Based on these two equations, we can easily obtain following equation

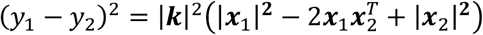

where, both |***x***_1_|^2^ and |***x***_2_|^2^ are equal to the length of the sequence. If the difference between two synonymous motifs is very great, ***x***_1_,***x***_2_ matched by these two motifs respectively should share much less bases 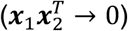. For an extreme case, the two sequence are totally different 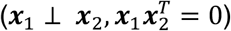. Based on the condition above including 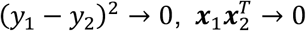 and the constant value of |***x***_1_|^**2**^ + |***x***_2_|^**2**^, we can infer that |***k***|^2^ → 0. Thus, the maximum activation value follows 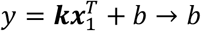. The AN becomes a dead AN if *b* ≤ 0. So, compared with the AN of SM that cannot represent two synonymous motif, the AN of MM representing the mixture of synonymous motifs exhibits a lower maximum activation value and a smaller weight. The importance of this kind of ANs for downstream ANs will be suppressed.

In a DCNN without the pooling layer, we further investigated an AN representing two synonymous motifs in deeper convolutional layers *i.* We assumed that there are no AN representing the mixture of synonymous motifs in layer 1 to layer *i* – 1. Based on this assumption, the feature map 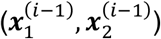 of ***x***_1_,***x***_2_ at layer *i* – 1 are of great difference especially for the key features with high activation values. The negative values of the feature map are set 0 by ReLU activation function 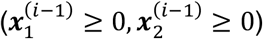. The key feature in 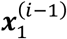 with high activation may be low activated or 0 in 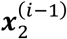 (for key feature *j*, 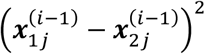 will be larger compared to the value of similar sequences).

It indicated that we were able to distinguish the sequences matched to the two synonymous motifs with the feature map of layer *i* – 1. The activation of the AN in layer *i* is the linear combination of the previous layer feature map (*y* = *f*(***x***) = *g*(***x***^(*i*–1))^ = ***k***(***x***^*i*–1^)^*T*^ + *b*). Similarly, for an AN in layer *i*, we can obtain the following equation

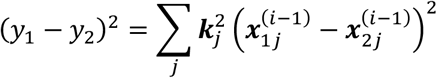

Where *j* is the feature number in layer *i* – 1. If this AN mixed the signals of 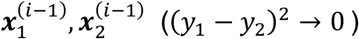, the result is the same with the first layer 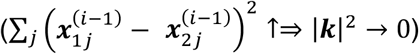.

However, the AN representing the mixture of synonymous motifs is usually accompanied by representing the strong consistent main motif (Fig. 2e). In layer *i* – 1 of this AN, for feature *s* representing the strong consistent main motif and feature *j* representing synonymous motifs, they may satisfy 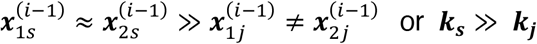. Although |***k***|^2^ of this kind of AN is smaller compared with the AN cannot recognize synonymous motifs, it can still obtain similar activation value *y* = ***k***(***x***^*i*–1)^)^*T*^ + *b* in two ways rather than becoming a dead kernel. One way is increasing 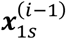 and 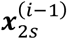 through the weight ***k***^(*i*–2)^ of previous layer neurons 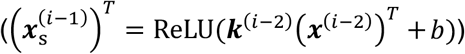. The other way is increasing ***k_s_*** and decreasing ***k_j_***, which may greatly reduce the side-effect of mixture of synonymous motifs. In a well-trained model, for an AN, compared to the large weight on the high activations of subsequence matched by the main motif, the signal generated by the subsequence matched by synonymous motif can be neglected. Otherwise, AN only representing strong synonymous motifs will destroy the robustness of the AN (Fig. 2e).

### Weighted sampling algorithm adapted from the genetic algorithm

The sequence sampling process is necessary to estimate the AN motifs. The first operation is the initialization of seed sequences. We randomly generated 5000 seed sequences that match the receptive field size. For each sequence, we randomly replaced a specific sub-sequence with one motif sequence of ANs in the previous layer. The position and the previous layer AN were randomly selected based on value of the normalized maximum contribution score:

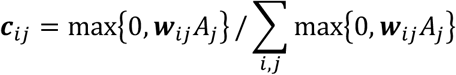

where *i* is the position number, *j* is the previous layer AN number and *A_j_* is the maximum activation value of the previous layer AN *j*. The second operation is sequence optimization. The sequence (*x*) is discrete so we cannot use the gradient decent method directly, so we adopt and adjusted the genetic algorithm. In one generation, we used the normalized gradient value *g* = *∂f*(***x***)/*∂****x*** as the probability to guide randomly select better mutation bases:

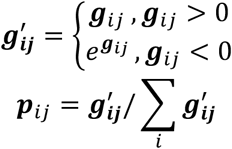

where *i* is the base of A,C,G,T, and *j* is the position. We kept 10% samples with top activation values in each generation. We randomly shifted 20% sequence samples based on the DCNN structure. The remaining samples were generated by roulette wheel selection and crossover operation. The total number of sequences did not change in each generation. The optimization would not stop until the maximum record of the mean activation value of each generation did not increase for 10 iterations. The third operation is sampling. At the end of each iteration in the genetic algorithm, sequence with positive activations were collected as samples. The duplicated sequences were removed. Based on the maximum activation value of existing samples, we split the activation value interval into 20 bins. We kept the number of samples in each bin less than 5000. If it was overflowed, we randomly selected 5000 samples among them. See Supplementary Information for the pseudo code of this algorithm.

### Shifting latent variable discovery and decoupling algorithm

For one AN, we need to design an algorithm to split the sample set according to the latent variables depending on the DCNN structure. From deep layers to shallow layers in DCNN model, when the result of a convolutional layer was the input of a max-pooling layer, the algorithm calculated the feature map of the convolutional layer and used K-means (K is the max-pooling size) to cluster the sequence samples into K subsets according to the features in feature maps. The algorithm would continuously cluster and split each subset reclusively once it found the result of convolutional layer was the input of the max-pooling layer. Finally, the number of subsets is 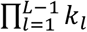 where *L* is the layer number of the AN and *k_l_* is the pooling size of the pooling operation applied on each convolutional layer. Based on each subset, we obtained the numerical estimation of PPMs. The algorithm can be applied on the newly generated subset again for decoupling the secondary important shifting motif if the samples are enough. This process has been shown in Fig. 2d and 2e. See Supplementary Information for details and the pseudo code of this algorithm.

### Algorithm implements

NeuronMotif were implemented in python. It depends on tensorflow and keras packages. Current version of NeuronMotif can only be applied to the DCNN implemented by tensorflow or keras. We only implemented the CPU version of NeuronMotif so it does not depend on GPU. The scripts are parallelized and can be run across the nodes of the computing cluster. The memory consumption depends on the DCNN structure and the AN receptive field size. We run the program on 4 servers. Each server contains 2 CPUs with 28 cores (Intel E5-2680) and 128GB memory. For each DCNN model mentioned in this work, the program can finish the decoupling of all convolutional ANs in about 3 days.

### Rebuilding and decoupling the DeepSEA and Basset models

DeepSEA and Basset are both 3-convolutional-layer models implemented in Torch, which is not compatible to NeuronMotif. We rewrote these two models with tensorflow and keras. We tried to keep the architecture, regularization, optimizer and so on consistent with the previous studies. We trained the models by the datasets that were used for training the original models. The dataset was split into a training set, a validation set and a test set according to the original papers. We also followed the training strategy described in the papers. We trained these two models with a single Nvidia P100 GPU card.

We applied NeuronMotif on DeepSEA and Basset. DeepSEA had (320, 480, 960) kernels in each convolutional layer. The max-pooling size was 4 for every convolutional layer. Theoretically, we would obtain 320,480 × 4 = 1920,960 × 4× 4 = 15360 AN motifs for L1, L2 and L3. However, some of them were absent for the dead kernel or low information content motifs that should be excluded. Similarly, Basset had (300, 200, 200) kernels and its max-pooling sizes were (3,4,4). Theoretically we would obtain 300,200 × 3 = 600, 200 × 3 × 4 = 2400 motifs from the L1, L2 and L3 of Basset Model.

### Synonymous motif mixture detection and diagnosis

To estimate motifs from each sample subset, we calculated its maximum activation value and activation value of consensus sequence. The max activation value is obtained by feeding all sample sequences to the substructure of the AN. The consensus sequence was obtained from the PPM of the motif. For each position, the nucleotide base with the largest probability among 4 bases in PPM was selected as the nucleotide in the consensus sequence. We fed the consensus sequence to the substructure of the AN and got the activation value. For all motifs in the same layer of the DCNN model, we can draw a scatter plot to find if serious synonymous motif mixture exists. It can be diagnosed by observing if activation values of consensus sequences are deviated from corresponding maximum activation values. More low activation values of the consensus sequences indicate more synonymous motif mixture in this DCNN model.

### Problematic neuron analysis

We investigated some problematic AN in the Basset model to find which part of the discovered motif makes the consensus sequence not be able to activate the AN. This is caused by the inconsistent sub-sequences at the certain position of the various sampled sequences playing key role in activating the AN. So, the consensus sequence of the motif cannot represent these sampled sequences. We call these motifs and their consensus sequences to be inconsistent, otherwise we call them to be consistent. Among the motif for an AN, consensus sequences of some motifs are consistent, which can activate the AN, but the inconsistent ones can’t. It is difficult to distinguish them by naked eyes because the information contents at different position are almost the same.

Here, we took one inconsistent and one consistent motif consensus sequence as examples. We aligned two sequences and used a 51bp window to slide on it. For each position, we replaced the sub-sequence (51bp) of the inconsistent consensus sequence with the corresponding sub-sequence of consensus sequence that can activate the AN to test if it can activate the AN. We found the valid position and tried to find the latent variables through clustering the sub-sequence-related feature maps or sub-sequence one-hot code that make it becomes mixture. Finally, we found they were the mixture of different synonymous motifs rather than the same shifted motif. See Supplementary Information for details.

### DCNN architecture optimization and deeper DCNN model construction

We tried to optimize the architecture solely without using regularization methods. Following the strategy of small kernels and max-pooling sizes, the kernel size and maxpooling size were set 3 and 2 respectively. We used ReLU as the activation function for each layer except for the last fully-connected layer with the sigmoid function. For Basset, we built a 5-convolutional-layer model BD-5 (kernel number and pooling operation: 32, pooling, 64, pooling, 128, pooling, 256, pooling, 512) and a 10-convolutional-layer model BD-10 (kernel number and pooling operation: 64, 64, pooling, 128, 128, pooling, 256, 256, pooling, 384, 384, pooling, 512, 512). At the end of convolutional layer, two fully-connected layers with 1024 and 164 ANs were appended. The number of kernel sizes was doubled based on the previous layer because the receptive field size was doubled for the deeper AN. In a longer receptive field, more combinations of the motifs need to be represented. For DeepSEA, we built a 10-convolutional layer model DD-10 (kernel number and pooling operation: 128, 128, pooling, 160, 160, pooling, 256, 320, pooling, 512, 640, pooling, 1024, 1280). At the end of convolutional layer, two fully-connected layers with 925 and 919 ANs were appended. However, the prediction performance of DD-10 was similar to DeepSEA. We found that the overlap of the first-layer receptive field is very small for the AN of the second layer. If we set the kernel size 3 in the first layer (receptive field size is 3bp), then the overlap proportion of the adjacent 3 ANs is 1/5 (receptive field size is 5bp). We need to get longer overlap by extending the kernel size in the first layer. We tried to train DD-10 with the first-layer kernel size equal to 5 (overlap proportion: 3/7 ≈ 43%), 7 (overlap proportion: 5/9 ≈ 56%) and 9 (overlap proportion: 7/11 ≈ 63%). The best one is the model with kernel size equal to 7 in the first layer. This result also matched the top convolution-pooling model in the ImageNet competition^25^. It seems to be a trade-off for the first kernel size. If it is too small, the structure is not good for training the second layer. On the contrary, the structure is not good for training the first layer. Hence, we finally set first-layer kernel size as 7 for the DD-10 and BD-10 model.

### Prediction performance comparison

Five models were involved in this work. They are Basset, BD-5, BD-10, DeepSEA and DD-5. After they had been trained on the training set and the validation set, they were tested on the test set. For each prediction target, we calculated the value of Area Under the Precision-Recall Curve (AUPRC). We used AUPRC rather than Area Under the Receiver Operating Characteristic curve (AUROC) because AUPRC is more sensitive to the unbalanced data. In the dataset of DeepSEA and Basset, the negative samples were much more than the positive samples so AUPRC is a better indicator. To compare and test the performance difference between models, we assume that if the performance of two model is the same, the difference of AUPRC value of the same prediction target is ΔAUPRC ∼ *N*(0, *σ*^2^). We did one-side *t*-test for each pair of models for comparison.

### Motif discovery

For each decoupled motif represented by the AN, we needed to filter and slice the motifs for regulatory elements. The decoupled motifs were generated by a sequence set. When the number of the sequences is very small, the motif is not reliable. We first applied the Laplace smoothing method to the PPMs of decoupled motifs. The smoothed PPM (**PPM**’) can be obtain by

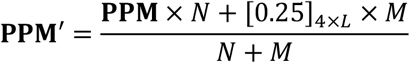

where *N* is the number of sequences that generate the PPM, [0.25]_4x*L*_ is the 4 × *L* matrix with all elements of 0.25, and *M* is the smoothing parameter. A larger *M* means a stronger smoothing process. We set *M* = 80 in our work. We regarded the nucleotide base position as a part of motif regions if its information content is greater than 1. We extended these motif regions with 3 bp at both the upstream and downstream. We merged these regions if they were overlapped. Regions longer than 8bp were regarded as motifs. We sliced these regions of PPM as the final discovered motifs. A large portion of these motifs can be matched with motifs in the JASPAR database. We showed a small portion of motifs in Fig. 5d with the motifStack^31^ package.

### Motif syntax discovery and validation

We used the ANs of layer 10 in BD-10 and DD-10 for the motif syntax discovery. We applied the decoupling algorithm twice for each AN and obtain 256 decoupled motifs. These decoupled motifs of the same AN usually shared similar shifted motifs. For convenient, we summarized the motifs by using Tomtom to match them to motifs in JASPAR. Based on the summarized TF motif set, we knew the arrangement of these TF motif. For instance, in Fig. 6a, the TF motif set includes CTCF and DDIT3:CEBPA and the arrangement of this two motif is CTCF-6N-DDIT3:CEBPA-[18-28N]-DDIT3:CEBPA-6N-CTCF, which is the motif syntax of the AN.

Except for literature validations such as JASPAR databases or some published papers, we also used ATAC-seq data to valid the motif syntax. If the motif syntax is real on genome, the region matched by the motif syntax should interact with some important molecules like TFs. Thus, the Tn5 transposes cutting frequency in the aligned regions may show footprinting. We collected five ATAC-seq datasets of five cell types or tissue including GM12878^2^, H1^32^, K562^33^, LNCaP^34^ and prostate^35^ (GSM1155957, GSM2264819, GSM2902637, GSM3632983, GSM3320984). We used the esATAC^36^ package developed by us to preprocess the dataset. For a concerned AN, we used it to scan the test data of Basset or DeepSEA. We collected the top 3000 activated regions, extended the regions to 1000bp and stack their Tn5 cutting frequency. We also counted the hard motif matching frequency at each position of these 1000bp regions with motifmatchr^37^.

### Code and more relative results

NeuronMotif code will be available at: https://github.com/wzthu/NeuronMotif

Relative results will be exibit at: https://wzthu.github.io/NeuronMotif

## Notes

https://wzthu.github.io/NeuronMotif/

## References

1 Searls, D. B. The language of genes. Nature 420, 211–217, doi:10.1038/nature01255 (2002).

2 Buenrostro, J. D., Giresi, P. G., Zaba, L. C., Chang, H. Y. & Greenleaf, W. J. Transposition of native chromatin for fast and sensitive epigenomic profiling of open chromatin, DNA-binding proteins and nucleosome position. Nature Methods 10, 1213–1218, doi:10.1038/Nmeth.2688 (2013).

3 Eraslan, G., Avsec, Z., Gagneur, J. & Theis, F. J. Deep learning: new computational modelling techniques for genomics. Nat Rev Genet 20, 389–403, doi:10.1038/s41576-019-0122-6 (2019).

4 Zhou, J. & Troyanskaya, O. G. Predicting effects of noncoding variants with deep learningbased sequence model. Nature Methods 12, 931–934, doi:10.1038/Nmeth.3547 (2015).

5 Kelley, D. R., Snoek, J. & Rinn, J. L. Basset: learning the regulatory code of the accessible genome with deep convolutional neural networks. Genome Research 26, 990–999, doi:10.1101/gr.200535.115 (2016).

6 Spitz, F & Furlong, E. E. M. Transcription factors: from enhancer binding to developmental control. Nat Rev Genet 13, 613–626 (2012).

7 Shrikumar, A., Greenside, P. & Kundaje, A. Learning important features through propagating activation differences. arXiv preprint arXiv:1704.02685 (2017).

8 Alipanahi, B., Delong, A., Weirauch, M. T. & Frey, B. J. Predicting the sequence specificities of DNA- and RNA-binding proteins by deep learning. Nature Biotechnology 33, 831–838, doi:10.1038/nbt.3300 (2015).

9 Searls, D. B. The Linguistics of DNA. Am Sci 80, 579–591 (1992).

10 Simonyan, K. & Zisserman, A. Very deep convolutional networks for large-scale image recognition. arXiv preprint arXiv:1409.1556 (2014).

11 Zou, J. et al. A primer on deep learning in genomics. Nature Genetics 51, 12–18 (2019).

12 He, Y., Shen, Z., Zhang, Q., Wang, S. & Huang, D.-S. A survey on deep learning in DNA/RNA motif mining. Briefings in Bioinformatics, doi:10.1093/bib/bbaa229 (2020).

13 Nguyen, A., Yosinski, J. & Clune, J. Multifaceted feature visualization: Uncovering the different types of features learned by each neuron in deep neural networks. arXiv preprint arXiv:1602.03616 (2016).

14 Koo, P. K. & Eddy, S. R. Representation learning of genomic sequence motifs with convolutional neural networks. Plos Computational Biology 15, doi:10.1371/journal.pcbi.1007560 (2019).

15 Jaganathan, K. et al. Predicting Splicing from Primary Sequence with Deep Learning. Cell 176, 535–548 e524, doi:10.1016/j.cell.2018.12.015 (2019).

16 Bogard, N., Linder, J., Rosenberg, A. B. & Seelig, G. A Deep Neural Network for Predicting and Engineering Alternative Polyadenylation. Cell 178, 91–106 e123, doi:10.1016/j.cell.2019.04.046 (2019).

17 Stormo, G. D. Introduction to protein-DNA interactions: structure, thermodynamics, and bioinformatics. (Cold Spring Harbor Laboratory Press, 2013).

18 Jolma, A. et al. DNA-Binding Specificities of Human Transcription Factors. Cell 152, 327–339, doi:10.1016/j.cell.2012.12.009 (2013).

19 Goodfellow, I. J., Shlens, J. & Szegedy, C. Explaining and harnessing adversarial examples. arXiv preprint arXiv:1412.6572 (2014).

20 Simonyan, K., Vedaldi, A. & Zisserman, A. Deep inside convolutional networks: Visualising image classification models and saliency maps. arXiv preprint arXiv:1312.6034 (2013).

21 Fornes, O. et al. JASPAR 2020: update of the open-access database of transcription factor binding profiles. Nucleic Acids Res 48, D87–D92 (2020).

22 Klemm, S. L., Shipony, Z. & Greenleaf, W. J. Chromatin accessibility and the regulatory epigenome. Nat Rev Genet 20, 207–220, doi:10.1038/s41576-018-0089-8 (2019).

23 Schwalie, P. C. et al. Co-binding by YY1 identifies the transcriptionally active, highly conserved set of CTCF-bound regions in primate genomes. Genome biology 14, doi: 10.1186/gb-2013-14-12-r148 (2013).

24 Molla, M., Delcher, A., Sunyaev, S., Cantor, C. & Kasif, S. Triplet repeat length bias and variation in the human transcriptome. Proceedings of the National Academy of Sciences of the United States of America 106, 17095–17100, doi:10.1073/pnas.0907112106 (2009).

25 Russakovsky, O. et al. ImageNet Large Scale Visual Recognition Challenge. Int J Comput Vision 115, 211–252, doi:10.1007/s11263-015-0816-y (2015).

26 Gupta, S., Stamatoyannopoulos, J. A., Bailey, T. L. & Noble, W. S. Quantifying similarity between motifs. Genome biology 8, doi:10.1186/gb-2007-8-2-r24 (2007).

27 Jolma, A. et al. DNA-dependent formation of transcription factor pairs alters their binding specificity. Nature 527, 384–388, doi:10.1038/nature15518 (2015).

28 Pugacheva, E. M. et al. Comparative analyses of CTCF and BORIS occupancies uncover two distinct classes of CTCF binding genomic regions. Genome biology 16, doi:10.1186/s13059-015-0736-8 (2015).

29 Grabowska, M. M. et al. NFI Transcription Factors Interact with FOXA1 to Regulate Prostate-Specific Gene Expression. Mol Endocrinol 28, 949–964, doi:10.1210/me.2013-1213 (2014).

30 Stormo, G. D. & Zhao, Y. Determining the specificity of protein-DNA interactions. Nat Rev Genet 11, 751–760, doi:10.1038/nrg2845 (2010).

31 Ou, J. H., Wolfe, S. A., Brodsky, M. H. & Zhu, L. H. J. motifStack for the analysis of transcription factor binding site evolution. Nature Methods 15, 8–9, doi:10.1038/nmeth.4555 (2018).

32 Liu, Q. et al. Genome-Wide Temporal Profiling of Transcriptome and Open Chromatin of Early Cardiomyocyte Differentiation Derived From hiPSCs and hESCs. Circ Res 121, 376–391, doi:10.1161/Circresaha.116.310456 (2017).

33 Calviello, A. K., Hirsekorn, A., Wurmus, R., Yusuf, D. & Ohler, U. Reproducible inference of transcription factor footprints in ATAC-seq and DNase-seq datasets using protocol-specific bias modeling. Genome biology 20, doi:10.1186/s13059-019-1654-y (2019).

34 Zhang, Z. D. et al. Loss of CHD1 Promotes Heterogeneous Mechanisms of Resistance to AR-Targeted Therapy via Chromatin Dysregulation. Cancer Cell 37, 584–598 e511, doi:10.1016/j.ccell.2020.03.001 (2020).

35 Park, J. W. et al. Reprogramming normal human epithelial tissues to a common, lethal neuroendocrine cancer lineage. Science 362, 91–95, doi:10.1126/science.aat5749 (2018).

36 Wei, Z., Zhang, W., Fang, H., Li, Y. D. & Wang, X. W. esATAC: an easy-to-use systematic pipeline for ATAC-seq data analysis. Bioinformatics 34, 2664–2665, doi:10.1093/bioinformatics/bty141 (2018).

37 Schep, A. N., Wu, B. J., Buenrostro, J. D. & Greenleaf, W. J. chromVAR: inferring transcription-factor-associated accessibility from single-cell epigenomic data. Nature Methods 14, 975–978, doi:10.1038/Nmeth.4401 (2017).

